# Kynurenine 3-monooxygenase (KMO) is a critical regulator of renal ischemia-reperfusion injury

**DOI:** 10.1101/272765

**Authors:** Xiaozhong Zheng, Ailiang Zhang, Margaret Binnie, Kris McGuire, Scott P Webster, Jeremy Hughes, Sarah E M Howie, Damian J Mole

## Abstract

Acute kidney injury (AKI) following ischemia-reperfusion injury (IRI) has a high mortality and lacks specific therapies. Here, we report that mice lacking kynurenine 3-monooxygenase (KMO) activity (*Kmo*^null^ mice) are protected against AKI after renal IRI. This advances our previous work showing that KMO blockade protects against acute lung injury and AKI in experimental multiple organ failure caused by acute pancreatitis. We show that KMO is highly expressed in the kidney and exerts major metabolic control over the biologically-active kynurenine metabolites 3-hydroxykynurenine, kynurenic acid and downstream metabolites. In experimental AKI induced by unilateral kidney IRI, *Kmo*^null^ mice had preserved renal function, reduced renal tubular cell injury, and fewer infiltrating neutrophils compared to wild-type (*Kmo*^wt^) control mice. Together, these data confirm that flux through KMO contributes to AKI after IRI, and supports the rationale for KMO inhibition as a therapeutic strategy to protect against AKI during critical illness.

## INTRODUCTION

In eukaryotes, the metabolic fate of the essential amino acid tryptophan is conversion via the kynurenine pathway into a range of metabolites that includes kynurenic acid, 3-hydroxykynurenine, and quinolinic acid. Enzymes involved in the metabolism of tryptophan along the kynurenine pathway are located throughout the body and brain, and are most abundant in the liver and kidney. The conversion of tryptophan to N-formylkynurenine (KYN) is catalyzed by tryptophan 2,3-dioxygenase (TDO) and indoleamine 2,3-dioxygenases (IDOs). The kynurenine pathway diverges at kynurenine into two distinct branches that are regulated by kynurenine aminotransferases (KATs) and kynurenine 3-monooxygenase (KMO) respectively (Figure 1). KMO is the only route of 3-hydroxykynurenine production known to occur in humans. KMO localizes to the outer membrane of mitochondria, and is highly expressed in peripheral tissues, including liver and kidney (Giorgini et al., 2013). KMO expression in kidney is mainly in tubular epithelial cells (60-80%) at both RNA and protein level in human (https://www.proteinatlas.org). 3-hydroxykynurenine is injurious to several cell types (Nakagami et al., 1996), causing tissue injury via oxidative stress, pathological cross-linking of proteins (Mizdrak et al., 2008) and inducing apoptotic cell death (Wang et al., 2014; Wilson et al., 2016). Kynurenine may also be metabolized to kynurenic acid by KATs and to anthranilic acid by kynureninase. Kynurenic acid is sedative (Harris et al., 1998) and has been shown to be protective against cell injury in certain inflammatory situations (Beninger et al., 1986; Vecsei et al., 2013).

**Figure 1.**
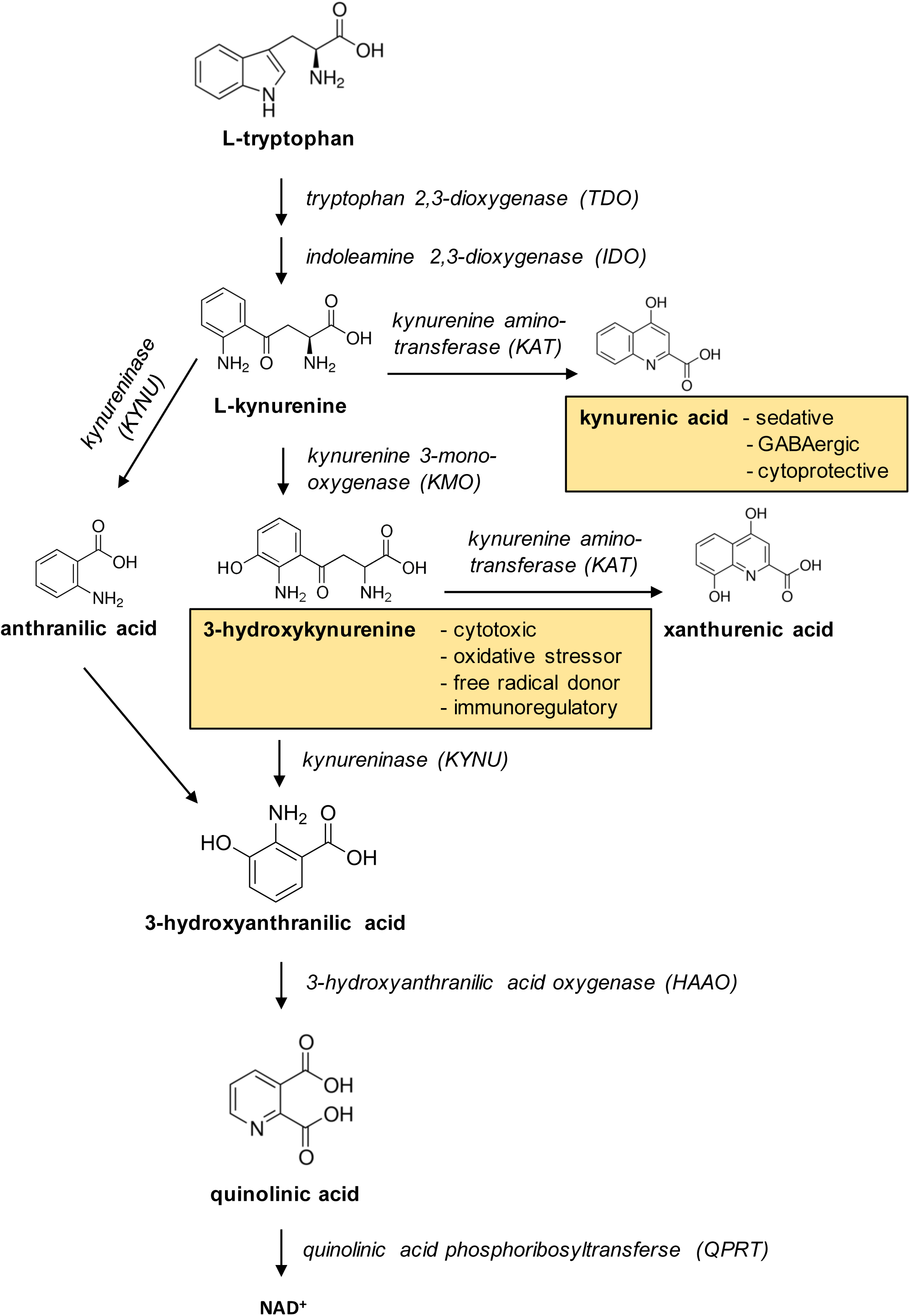
Overview of the kynurenine pathway of tryptophan metabolism. 3-hydroxykynurenine, the product of the gate-keeper enzyme kynurenine 3-monooxygenase (KMO), and kynurenic acid, one of the other branch motablite are highlighted.

Renal ischemia-reperfusion injury (IRI) is a leading cause of acute kidney injury (AKI). AKI, as a component of multiple organ dysfunction syndrome (MODS), has a high mortality and lacks specific therapies. We have recently demonstrated that increased flux through KMO is associated with the severity of critical illness in human patients with MODS triggered by acute pancreatitis (AP), and plasma concentrations of 3-hydroxykynurenine correlate with the burden of inflammation (Skouras et al., 2016). In AP-MODS, renal dysfunction is second only to respiratory dysfunction in incidence (Mole et al., 2009). In experimental rodent models of AP-MODS, in which AKI is an important contributor, we have shown that pharmacological inhibition of KMO, and separately, transcriptional blockade of the *Kmo* gene, reduces 3-hydroxykynurenine formation and protects against lung and kidney injury. The exact mechanisms that drive AKI in AP are not well understood, but are likely to stem from a combination of hypoperfusion-reperfusion (IRI) and metabolic toxicity.

KMO blockade through systemic administration of a KMO inhibitor protects against AKI during experimental MODS in rats but it is not known whether this is due to direct inhibition of KMO in kidney tissue itself, or a secondary effect of generalised KMO blockade. To further elucidate the role of renal KMO in protecting against experimental AKI, we tested whether kidneys in *Kmo*^null^ mice are protected from experimental AKI induced by the direct insult of renal IRI.

## RESULTS

### Kmo^null^ mice are protected against acute kidney injury after renal IRI

To determine whether absent KMO activity affected the severity of AKI after IRI, we compared the effect of experimental IRI in *Kmo*^wt^ and *Kmo*^null^ mice. *Kmo*^null^ mice were protected against the effects of renal IRI, experiencing less severe AKI, demonstrated by a lower plasma creatinine and less tubular damage than *Kmo*^wt^ control mice (Figure 2). The experimental model of IRI was effective at inducing severe AKI, because IRI caused an elevated plasma creatinine to 190.5 ± 26.3 µmol/L at 24 hours after IRI in *Kmo*^wt^ mice, compared to a plasma creatinine of 23.7 ± 5.5 µmol/L at the equivalent time after sham operation in *Kmo*^wt^ mice. *Kmo*^null^ mice had a lower plasma creatinine 24 hours after IRI (111.2 ± 21.3 µmol/l) compared to the equivalent value in *Kmo*^wt^ control mice. This difference was statistically significant (P<0.05) (Figure 2a) with a magnitude that is biologically relevant. Renal IRI also significantly elevated the urinary albumin/creatinine ratio (ACR) to 2590 ± 871.5 µmol/L at 24 hours after IRI in *Kmo*^wt^ mice, compared to ACR of 19.7 ± 5.0 µmol/L at the equivalent time after sham operation in *Kmo*^wt^ mice (P<0.01). Urinary albumin/creatinine ratio was profoundly increased after IRI both in *Kmo*^wt^ and *Kmo*^null^, confirming AKI, but the magnitude of this elevation in *Kmo*^null^ is smaller than that in *Kmo*^wt^ mice although this difference is not statistically significant (P>0.999, Figure 2b). Because a previous report of a different strain of KMO-deficient mice (Korstanje et al., 2016) reported proteinuria in unstressed KMO-deficient mice, we measured plasma and urine albumin concentrations (Supplementary Figure 1a and 1b). Contrary to that previous report, urine albumin concentrations were not different between sham-operated *Kmo*^null^ and *Kmo*^wt^ mice, and specifically not lower in *Kmo*^null^ mice.

**Figure 2.**
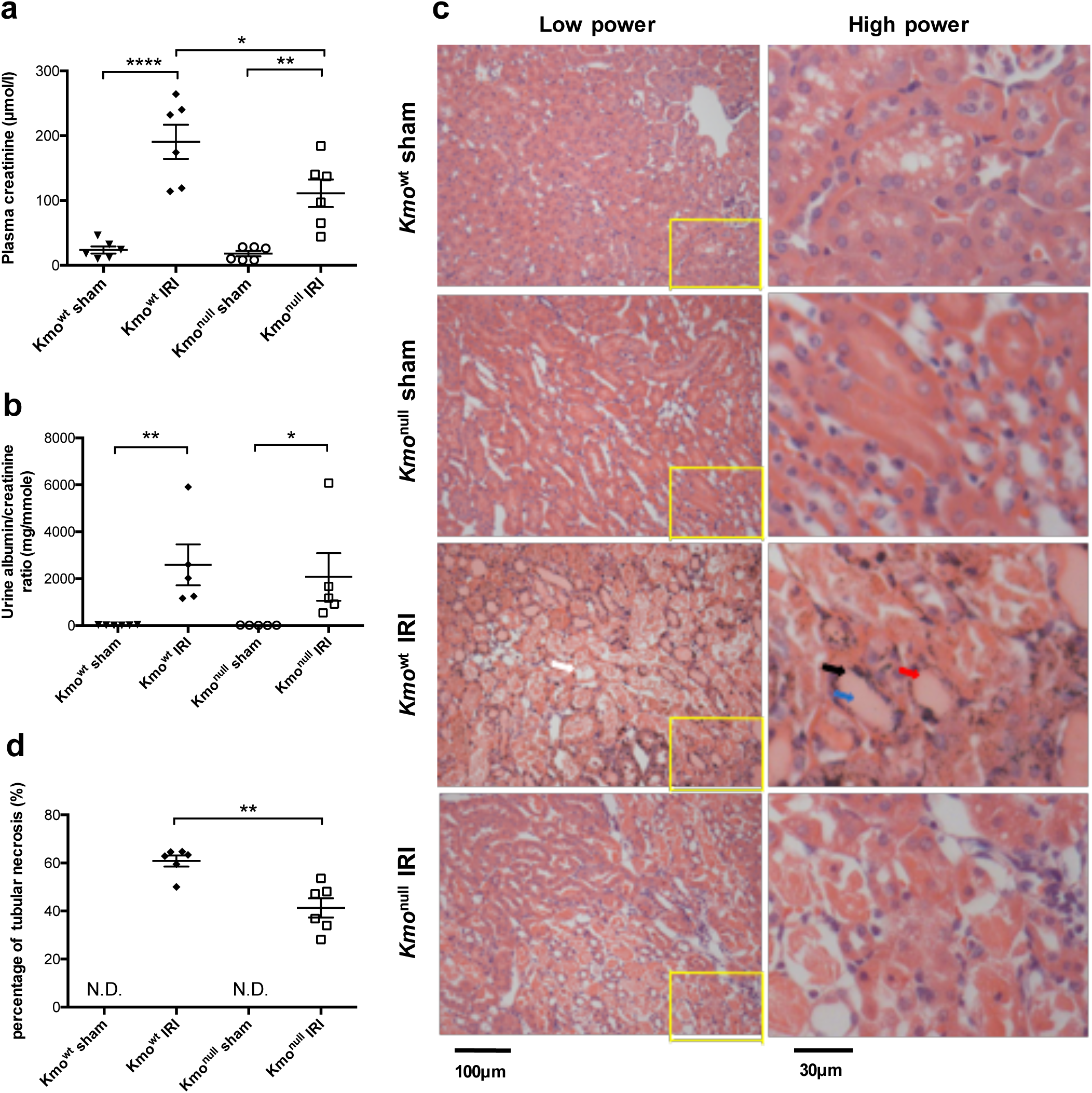
Biochemical indices of renal function and histological assessment of renal tubule injury after ischemia-reperfusion injury (IRI) in *Kmo*^wt^ and *Kmo*^null^ mice. (**a**) Plasma creatinine. (**b**) Urine albumin/creatinine ratio. (**c**) Representative digital micrographs of histological sections of formalin-fixed paraffin-embedded kidney tissue from *Kmo*^wt^ and *Kmo*^null^ mice after IRI or sham operation, stained with haematoxylin and eosin and visualized by light microscopy at ×200 original magnification. Low power images are showed on the left panel. The relative image of selected area are showed in high power on the right pannel. Tubular necrosis (white arrow), loss of the brush border (black arrow), cast formation (blue arrow), and tubular dilatation (red arrow) are indicated within the outer stripe of the renal medulla. (**d**) Enumeration of necrotic renal tubules expressed as a percentage of all tubules. N.D: not detected. **For all panels,** mice were subjected to unilateral nephrectomy and contralateral IRI, or sham operation, under general anaesthesia as described. 24 hours after IRI or sham operation, mice were euthanased and blood, urine and kidney sampled for analysis. All graphs show data from individual mice (one data point per mouse), with lines showing mean ± SEM. Group sizes were n=6 mice per group, except for panel **b**, where urine was only successfully obtained from n=5 mice per group (individual data shown). Statistically-significant differences between groups were analyzed by one-way ANOVA with post hoc Tukey’s test; * P<0.05; ** P<0.01 and **** P<0.0001.

When histological tubular damage was assessed, *Kmo*^null^ mice were protected from structural AKI after IRI, congruent with the functional protection indicated by the plasma creatinine concentrations. *Kmo*^wt^ mice sustained severe tubular damage after IRI, evidenced by widespread tubular necrosis, loss of the brush border, cast formation, and tubular dilatation within the outer stripe of the outer medulla (Figure 2c). On histological examination, kidneys from *Kmo*^null^ mice showed significantly less tubular damage after IRI compared to equivalent tissue sections from *Kmo*^wt^ mice. Kidneys from sham-operated mice from either *Kmo*^wt^ or *Kmo*^null^ strain incurred no visible tubular injury on histological assessment. Quantification of tissue injury, obtained by counting necrotic tubules expressed as a percentage of all tubules, was significantly lower in *Kmo*^wt^ IRI mice (41.3 ± 4.0 %) than *Kmo*^wt^ IRI mice (60.8 ± 2.3 %) (P<0.01, Figure 2d). Together, these data clearly show that absent KMO activity in kidney tissue leads to functional and histological renal protection from structural AKI following experimental IRI.

### Tubular epithelial cell apoptosis is reduced in the kidney of Kmo^null^ mice following IRI

Because therapeutic KMO inhibition protects against renal tubular cell apoptosis in AKI during experimental AP in rats, and because the KMO product 3-hydroxykynurenine induces apoptosis of cells in vitro, we examined the extent of tubular cell apoptosis. We labelled and enumerated apoptotic renal tubular cells using the terminal deoxynucleotidyl-transferase–mediated dUTP nick-end labelling (TUNEL) assay. There was no difference in the baseline apoptotic cell count in renal tubules in *Kmo*^wt^ and *Kmo*^null^ mice. Experimental IRI was a powerful inducer of apoptosis in renal tubular cells, causing substantial tubular epithelial cell apoptosis detectable at 24 hours after IRI (Figure 3a). Importantly, although *Kmo*^null^ mice sustained a degree of renal tubular cell apoptosis, the number of apoptotic renal tubular cells was significantly lower in *Kmo*^null^ mice with IRI (19.8 ± 4.5 /field) than in *Kmo*^wt^ control mice (49.9 ± 11.6 / field) (P<0.05, Figure 3b).

**Figure 3.**
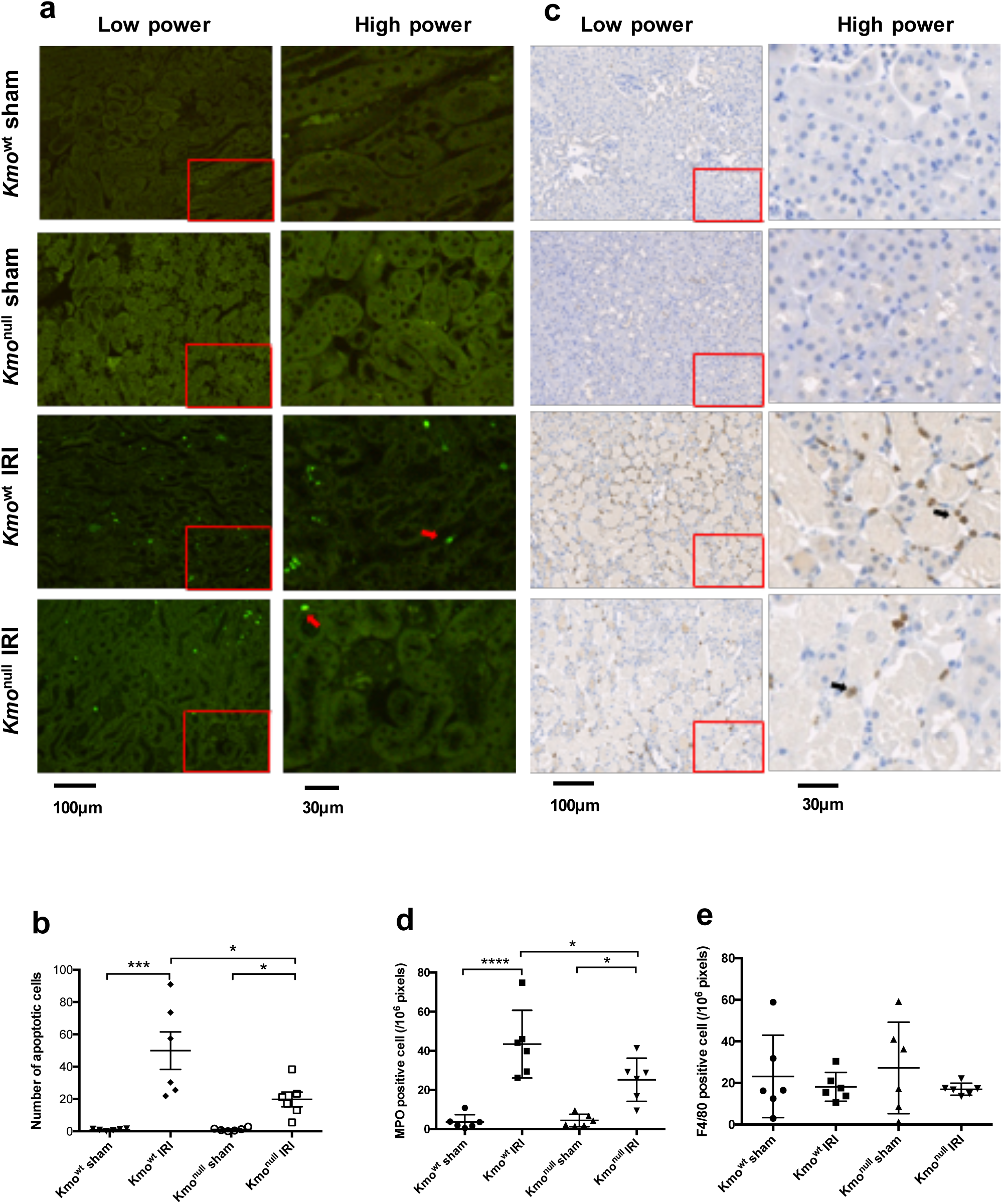
Tubular epithelial cell apoptosis, neutrophil infiltration and monocyte-derived macrophage accumulation in kidney tissue after IRI. (**a**) Representative digital micrographs of TUNEL-stained kidney sections at ×100 original magnification. Red arrows indicate TUNEL^+^ apoptotic cells. Low power images are showed on the left panel. The relative image of selected area are showed in high power on the right pannel. (**b**) Number of apoptotic cells per low-power field. (**c**) Representative digital micrographs selected from scanned MPO-stained entire kidney images. Black arrows indicate MPO^+^ cells. Low power images are showed on the left panel. The relative image of selected area are showed in high power on the right pannel. (**d**) MPO^+^ neutrophil infiltration, expressed as MPO^+^ cells per 10^6^ pixels. (**e**) F4/80^+^ monocyte-derived macrophage accumulation, expressed as F4/80^+^ cells per 10^6^ pixels**. For all panels,** *Kmo*^wt^ and *Kmo*^null^ mice were subjected to IRI or sham operation as described and euthanased 24 hours afterwards. Kidneys were sampled for analysis. Apoptotic cells were labelled by TdT-mediated dUTP nick-end labelling (TUNEL) assay and enumerated from digitally-scanned micrographs using ImageJ. Neutrophils and monocyte-derived macrophages were labelled by immunohistochemistry using antibodies to myeloperoxidase (MPO) and F4/80 respectively, visualized by diaminobenzoate and enumerated. All graphs show data from individual mice (one data point per mouse), with lines showing mean ± SEM. Group sizes were n=6 mice per group. Statistically-significant differences between groups were analyzed by one-way ANOVA with post hoc Tukey’s test; * P<0.05; and **** P<0.0001.

### KMO deletion inhibits neutrophil infiltration in the kidney following IRI

Acute kidney injury incorporates an element of acute inflammation. Therefore, we directly measured neutrophil accumulation in the kidney following IRI by immunohistochemical staining for myeloperoxidase (MPO) and enumeration of MPO^+^ cells. A clearly measurable influx of MPO^+^ neutrophils was identified on sections of kidney tissue in both *Kmo*^wt^ and *Kmo*^null^ mice, 24 hours after IRI, compared to very low numbers of MPO^+^ cells in kidneys of sham-operated animals (Figure 3c). Using quantitative analysis, we detected a lower number of accumulated neutrophils in the kidneys of *Kmo*^null^ mice (25.2 ± 4.5 /10^6^ pixels) compared to *Kmo*^wt^ mice after IRI (43.4 ± 7.1 /10^6^ pixels), and this difference was statistically significant (P<0.05, Figure 3d). We also examined monocyte-derived macrophage accumulation after IRI by F4/80 immunohistochemistry. There was no significant infiltrate of cells positive for the mouse monocyte marker F4/80 in the kidney at 24 hours after renal IRI in either *Kmo*^null^ mice (17.0 ± 1.2 /10^6^ pixels) or *Kmo*^wt^ mice (18.2 ± 2.8 /10^6^ pixels) (Figure 3e).

### Renal IRI downregulates kynurenine pathway enzyme mRNA expression and upregulates interleukin-6 and tumour necrosis factor-α mRNA expression

Next, we asked whether IRI affected expression of kynurenine pathway enzymes, specifically kynurenine 3-monooxygenase (*Kmo*), kynureninase (*Kynu*), kynurenine amino-transferase (*Kat2*), 3-hydroxyanthranilic acid oxygenase (*Haao*) and Quinolinate phosphoribosyltransferase (*Qprt*) in kidney tissue. Using real time PCR, we detected a profound and statistically significant decrease in *Kmo* mRNA expression in kidney tissue after IRI in *Kmo*^wt^ mice compared to baseline expression levels in sham-operated *Kmo*^wt^ mice. There was no detectable *Kmo* mRNA expression in kidneys from *Kmo*^null^ mice, in keeping with the expected genotype and with our previous experience with this mouse strain (Figure 4a). Interestingly, IRI caused a reduction in mRNA expression in both *Kmo*^wt^ and *Kmo*^null^ mouse strains for *Kynu, Kat2, Haao* and *Qprt* when compared to sham-operated mice of both transgenic strains. There was no strain-specific difference in the expression of *Kynu, Kat2, Haao* and *Qprt* mRNA attributable to *Kmo* gene deletion in *Kmo*^null^ mice (Figure 4b, 4c, 4d and supplementary Figure 2). The observed reduction in kynurenine pathway enzyme mRNA expression cannot be explained by experimental artefact, because quantification of mRNA expression of the pro-inflammatory cytokines IL-6 and TNFα in the same cDNA preparations used for kynurenine pathway transcripts was significantly upregulated following IRI, and equally so in *Kmo*^wt^ and *Kmo*^null^ mouse strains. Sham-operated mouse kidneys from both *Kmo*^wt^ and *Kmo*^null^ strains had very low expression of IL-6 and TNFα mRNA. Mice subjected to IRI demonstrated strong up-regulation of IL-6 and TNFα compared to sham-operated mice: IL-6 mRNA expression in the IRI kidney increased 173 fold in *Kmo*^wt^ mice and 97.3 fold in *Kmo*^null^ mice, respectively (Figure 4e); TNFα mRNA expression in the IRI kidney increased 3.9 fold in *Kmo*^wt^ mice and 2.1 fold in *Kmo*^null^ mice (Figure 4f). The difference in IL-6 and TNFα mRNA upregulation in IRI was not statistically significant between *Kmo*^wt^ mice and *Kmo*^null^ mice. Together, these data show that renal IRI drives a downregulation of kynurenine pathway enzyme mRNA expression, or potentially reflects loss of tubular cells through necrosis, independent of the functionality of *Kmo*, and simultaneously upregulates pro-inflammatory cytokine expression.

**Figure 4.**
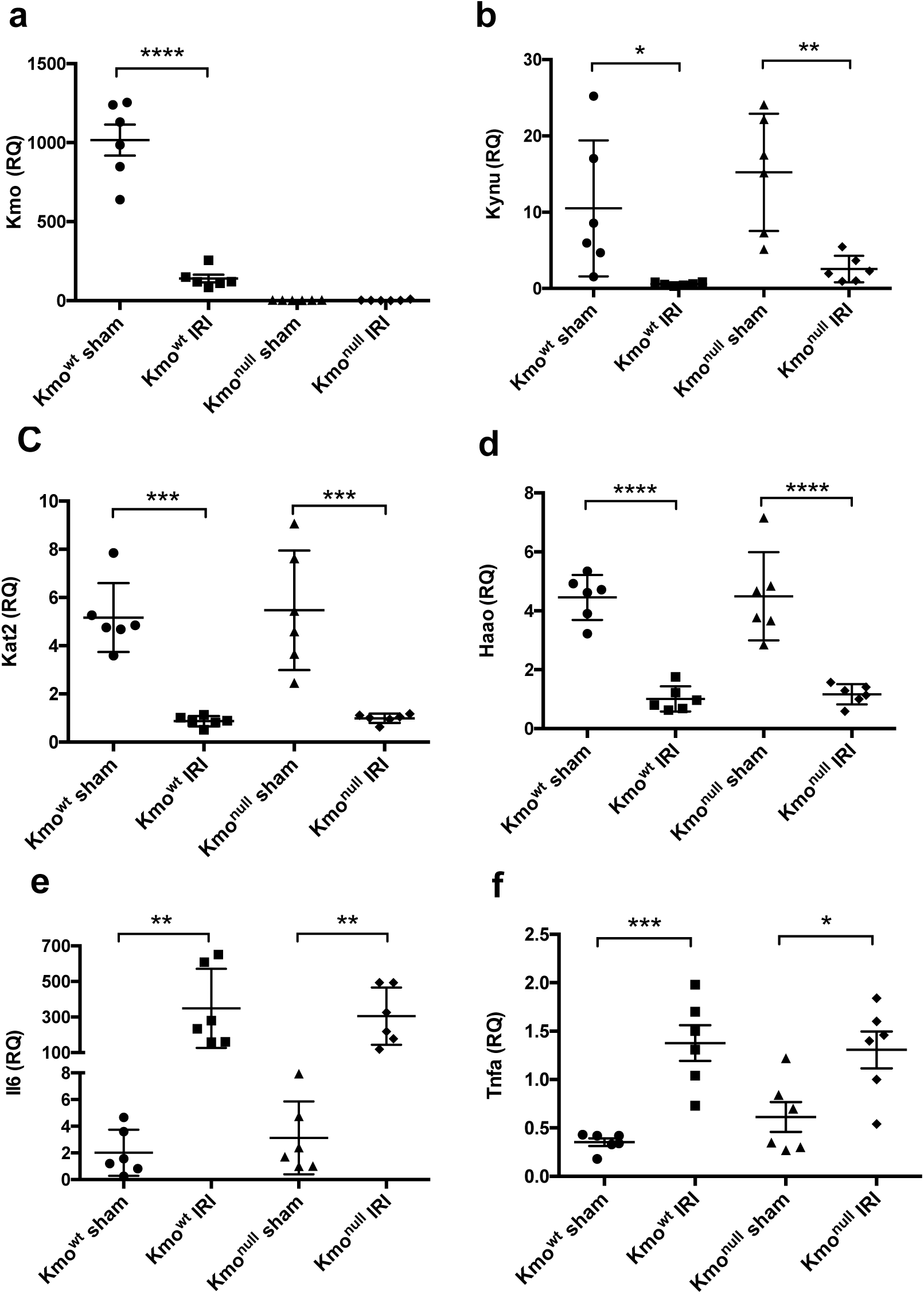
mRNA expression of kynurenine pathway enzymes and pro-inflammatory cytokines in kidney tissue after IRI. (**a**) kynurenine 3-monoxygenase, *Kmo*; (**b**) kynureninase, *Kynu*; (**c**) kynurenine aminotransferase, *Kat2*; (**d**) 3-hydroxyanthranilic acid oxidase, *Haao*; (**e**) interleukin-6, *Il6*; (**f**) tumour necrosis factor α, *Tnfa.* **For all panels**, *Kmo*^wt^ and *Kmo*^null^ mice were subjected to IRI or sham operation and euthanased 24 hours afterwards. Kidney tissue was sampled, snap frozen in liquid nitrogen and RNA subsequently extracted for analysis by real-time PCR as described. mRNA levels of the target gene were normalized to 18S ribosomal RNA and are presented as relative quantification (RQ) values. All graphs show data from individual mice with lines showing mean ± SEM. Group sizes were n=6 mice per group. Statistically-significant differences between groups were analysed by one-way ANOVA with post hoc Tukey’s test; * P<0.05; and **** P<0.0001.

### Kynurenine metabolite changes in experimental IRI in mice

The metabolic product of KMO, 3-hydroxykynurenine, is injurious to cells and tissues. Because *Kmo*^null^ mice are unable to form 3-hydroxykynurenine and are also protected from AKI, it was important to confirm absent 3-hydroxykynurenine in plasma and kidney tissue after IRI. Furthermore, because the biochemical phenotype of *Kmo*^null^ mice is a diversion of kynurenine metabolism to kynurenic acid, and because we had observed that IRI down regulates kynurenine pathway enzyme mRNA expression, we measured kynurenine metabolite concentrations in plasma and tissue (Figure 5). In sham-operated animals, we observed the plasma biochemical phenotype expected of *Kmo*^null^ mice, namely reduced (or absent) 3-hydroxykynurenine levels, reduced 3-hydroxyanthranilic acid levels, an upstream backlog of kynurenine, and a metabolic diversion of kynurenine to kynurenic acid. Tryptophan concentrations were not different between sham-operated *Kmo*^wt^ and *Kmo*^null^ mice. After IRI, the kynurenine pathway biochemical phenotype showed intriguing changes: plasma tryptophan was depleted. Tryptophan concentration in *Kmo*^null^ mice was significantly lower in the IRI group (5501 ± 999 ng/ml) than in sham operated group (9411.0 ± 588.2 ng/ml) (P<0.01). Tryptophan concentration in *Kmo*^wt^ mice was also significantly lower in the IRI group (3232 ± 98.6 ng/ml) than in sham operated group (8560 ± 893.5 ng/ml) (P<0.001). The degree of tryptophan depletion was less pronounced in *Kmo*^null^ mice than *Kmo*^wt^ mice after IRI, but this was not statistically significant between these two mouse strains (Figure 5a). Kynurenine concentrations were not altered in IRI to an extent that exceeded the profound differences at baseline between *Kmo*^wt^ and *Kmo*^null^ mouse strains (Figure 5b). Experimental IRI caused an elevation in the potentially protective molecule kynurenic acid that was significantly further increased in *Kmo*^null^ mice with IRI (*Kmo*^null^ IRI 8738.0 ± 673.6 ng/ml and *Kmo*^wt^ IRI 581.6 ± 72.9 ng/ml. P<0.0001, Figure 5c). Although IRI induced a rise in 3-hydroxyanthranilic acid, this was not different between *Kmo*^wt^ and *Kmo*^null^ mouse strains (*Kmo*^null^ IRI 324.3 ± 93.6 ng/ml and *Kmo*^wt^ IRI 389.0 ± 33.3 ng/ml. Figure 5d). Concentrations of 3-hydroxykynurenine were significantly increased 24 hours after IRI in *Kmo*^wt^ mice in plasma (IRI 53.3 ± 6.3 ng/ml and sham 34.0 ± 6.3 ng/ml. P<0.05, Figure 5e), but not in kidney tissue (Figure 4f), and, as expected, *Kmo*^null^ mice showed extremely low levels of 3-hydroxykynurenine both in plasma and kidney tissue (Figure 5e and 5f). Together, these data convincingly demonstrate that 3-hydroxykynurenine production by KMO is a significant contributor to AKI following renal IRI, and reinforce the concept of KMO inhibition as a protective strategy to protect against organ dysfunction in critical illness.

**Figure 5.**
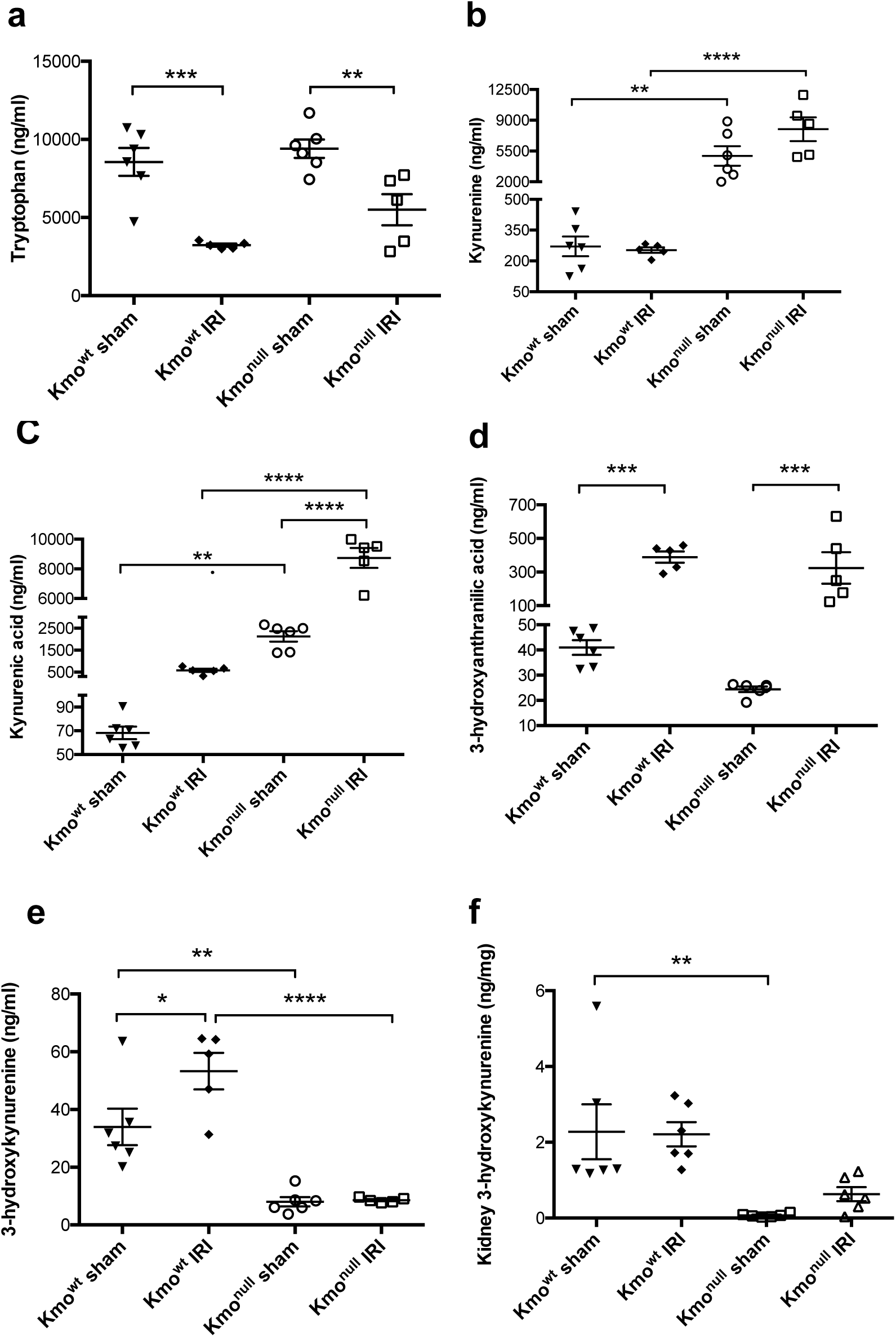
Kynurenine pathway metabolite concentrations in plasma and kidney tissue after IRI. (**a**) Plasma tryptophan; (**b**) plasma kynurenine; (c) plasma kynurenic acid; (d) plasma 3-hydroxyanthranilic acid; (e) plasma 3-hydroxykynurenine; (f) kidney tissue 3-hydroxykynurenine. **For all panels,** mice were subjected to unilateral nephrectomy and contralateral IRI, or sham operation, under general anaesthesia. 24 hours after IRI or sham operation, mice were euthanased and blood and kidney tissue sampled for analysis. Extracts of plasma (**panels a–e**) or kidney tissue (**panel f**) were analysed by LC-MS/MS as described. All graphs show data from individual mice (one data point per mouse), with lines showing mean ± SEM. Group sizes were n=6 or n=5 (where one plasma was not obtained) mice per group. Statistically-significant differences between groups were analysed by one-way ANOVA with post hoc Tukey’s test; * P<0.05; ** P<0.01 *** P<0.001 and **** P<0.0001.

## DISCUSSION

Kynurenine metabolites are generated by tryptophan catabolism and regulate biological processes that include host-microbiome signalling, immune cell response, and neuronal excitability (Cervenka et al., 2017). More recently, we and others are focusing on the role of the kynurenine pathway in inflammation and tissue injury. In particular, we have focused on the pathway enzyme kynurenine 3-monooxygenase (KMO), which plays a key gate-keeper role in kynurenine metabolism, by determining the metabolic fate of kynurenine generated by upstream dioxygenases. Recent genetic analysis in mice identified Kmo as a candidate gene associated with albuminuria. Glomerular KMO expression is decreased in both mouse and human kidneys in a diabetic environment (Korstanje et al., 2016). These previous reports support a link between KMO expression and healthy kidney function. In the present work, we conclusively demonstrate that KMO is a critical regulator of tissue injury in kidneys subjected to IRI. We used a transgenic mouse strain (*Kmo*^null^ mice) that lacks KMO activity in all tissues by disruption of transcription in exon 5 of the *Kmo* gene, and compared it to *Kmo*^wt^ mice with normal KMO activity. The key finding of our paper is that *Kmo*^null^ mice are significantly protected from AKI after IRI, measured by preserved biochemical renal function and histological evidence of protection against tubule cell necrosis and apoptosis. Furthermore, neutrophil infiltration was reduced in *Kmo*^null^ mice with IRI, when compared to *Kmo*^wt^ mice as controls.

Tryptophan metabolism through the kynurenine pathway is most widely studied to date in relation to disorders of the nervous system, for example Huntington’s disease, stress-related depression, schizophrenia, Alzheimer’s disease and Parkinson’s disease (Schwarcz and Stone, 2017). In recent years the regulation of kynurenine metabolism has been increasingly evaluated in relation to acute tissue injury, and has become an attractive therapeutic target in several disease areas (Thevandavakkam et al., 2010). Our data here show altered kynurenine metabolite levels in *Kmo*^null^ mice after IRI compared to *Kmo*^wt^ IRI mice. Despite showing comparable downregulated expression of the kynurenine pathway enzymes following experimental IRI, levels of kynurenine and kynurenic acid were significantly higher in *Kmo*^null^ mice compared to *Kmo*^wt^ controls. Whilst such changes are characteristic of the *Kmo*^null^ phenotype, kynurenine and kynureninic acid levels were further increased in *Kmo*^null^ IRI mice compared to *Kmo*^null^ sham-operated mice. These metabolites may contribute to protection from AKI after IRI in *Kmo*^null^ mice, particularly since kynurenic acid has demonstrated a protective role in a number of other inflammatory situations through its activity at glutamate receptors. Levels of free radical-generating 3-hydroxyanthranalic acid were increased to a similar degree in both mouse strains following induction of IRI suggesting that this metabolite is not implicated in the development of AKI. Since plasma 3-HK production was significantly higher in *Kmo*^wt^ IRI mice than in all other conditions, it seems likely that 3-HK-mediated effects contribute to the observed exacerbated response to IRI.This study provides further evidence that metabolic flux through the kynurenine pathway is upregulated in systemic inflammation, congruent with clinical observations in humans with AP (Mole et al., 2008; Skouras et al., 2016), trauma (Logters et al., 2009; Pellegrin et al., 2005), undergoing coronary artery bypass surgery (Forrest et al., 2011), sepsis (Wang et al., 2010), and chronic renal failure (Pawlak et al., 2009).

Plasma concentrations of 3-hydroxykynurenine correlate with the burden of inflammation, incidence of organ dysfunction and AP severity in human (Skouras et al., 2016). Altered kynurenine pathway metabolite levels observed here suggest that KMO also plays a critical role in kidney injury, strengthening the rationale for the use of systemic KMO inhibitors in this indication.

*Kmo*^null^ mice were generated and characterized in our laboratory to investigate the role of 3-hyxdroxykynurenine as an important effector of tissue injury. In *Kmo*^null^ mice, fecundity, fertility and longevity up to 2 years of age are not affected, from which we can infer that prolonged KMO blockade and consequent chronic exposure to significantly elevated concentrations of kynurenic acid and kynurenine is well tolerated, at least in adapted mice (Mole et al., 2016). Other researchers have also independently demonstrated that the deleted *Kmo* was not essential for embryonic and postnatal survival (Giorgini et al., 2013). KMO depletion decreased plasma 3HK level and increased plasma KA level, which may be potentially protective against renal damage caused by IRI.

Kidney ischemia–reperfusion injury is an inevitable consequence of the procedure of kidney transplantation and its severity has been correlated with an increased incidence of both acute and chronic rejection (Nankivell and Alexander, 2010; Wu et al., 2014). Apoptosis of tubular epithelial cells contributes to the development of ischemic AKI, and injury at this site contributes to organ failure. Inhibiting apoptosis, both before and after renal insults, will inevitably improve the outcome of human AKI (Havasi and Borkan, 2011). Neutrophils and monocytes/macrophages are important contributors to ischemic kidney injury and repair. Neutrophils attach to the activated endothelium and accumulate in the kidney both in animal models and in human AKI (Bonventre and Yang, 2011). Recent studies have shown that proinflammatory cytokines and chemokines such as TNF-α, IL-6, MCP-1, and MIP-2 contribute to the development of renal IRI (Zhou et al., 2017). In the present study, we report the original finding that genetic KMO deletion provides protection against kidney damage caused by IRI. Renal injury was observed on histological sections of kidney tissue from mice with IRI. They also showed an increase in neutrophilic inflammation and increased apoptosis. All these features of renal injury were essentially reduced in KMO depletion mice. Additionally, we observed that the mRNA expression of pro-inflammatory cytokines such as TNF-α and IL-6 increased in IRI mice compared to sham animals which has been reported earlier in human AP patients (Skouras et al., 2016). There was no significant difference about the TNFα and IL-6 mRNA expression between *Kmo*^null^ and *Kmo*^wt^ IRI mice.

In summary, our study shows *Kmo*^null^ mice had preserved renal function, reduced renal tubule cell injury and apoptosis, and fewer infiltrating neutrophils compared to *Kmo*^wt^ control mice. Together, these data strongly support the translational potential of KMO inhibition as a therapeutic strategy to protect against renal injury in acute inflammation.

## EXPERIMENTAL PROCEDURES

### Ethical considerations

All experiments were performed after Research Ethics Committee and Veterinary review at The University of Edinburgh, and were conducted according to the United Kingdom Use of Animals (Scientific Procedures) Act 1986, under license PPL60/4250.

### Animals

Mice on a C57BL/6 background engineered to lack KMO activity by insertion of a polyA transcription ‘stop’ motif before exon 5 of the *Kmo* gene (*Kmo*^*tm1a(KOMP)Wtsi*^), hereafter referred to as *Kmo*^null^ mice, were generated and maintained in our laboratory as previously described (Mole et al., 2016). Control mice (*Kmo*^*tm1c(KOMP)Wtsi/flox(ex5)*^), with normal *Kmo* gene transcription and KMO activity with *loxP*-flanked exon 5 of *Kmo* but no *cre*-recombinase expression and therefore a wild-type phenotype, hereafter referred to as *Kmo*^wt^ mice, were generated from the same founders as the *Kmo*^null^ strain. Genotyping of all mice was performed by polymerase chain reaction (PCR) using primers and a protocol published previously (Mole et al., 2016). Male mice only were used. Mice were 10–15 weeks old and were housed under specific pathogen-free conditions in the Biomedical Research Resources Facility of the University of Edinburgh.

### Experimental ischemia/reperfusion injury (IRI)

Experimental kidney IRI was induced as described previously (Hesketh et al., 2014). Briefly, mice were given a general anaesthetic using intraperitoneal ketamine and metomidate according to local dosage guidelines. Under aseptic conditions, a midline laparotomy and right nephrectomy were performed. The left renal pedicle was identified and occluded by atraumatic clamp for 22 minutes. The duration of clamping was determined by our previous experience with this model using the same background mouse strain. Body temperature was maintained at 35°C using a homeostatically-controlled blanket (Harvard Apparatus, Boston, MA). After reperfusion, the abdominal wall was sutured closed with 5-0 polypropylene and the skin closed with metal clips. Anaesthesia was reversed with Antisedan. Fluid resuscitation with 1 mL of sterile 0.9% NaCl was administered subcutaneously to the scruff after surgery. Sham-operated mice underwent general anesthesia, laparotomy and unilateral nephrectomy, but no clamping of the left renal pedicle. The animals were singly-housed and maintained in a 28°C warm box overnight to recover from surgery. After 24 hours, mice were humanely killed under terminal general anesthesia. Blood was collected into tubes containing EDTA (BD Biosciences). Urine was collected into sterile Eppendorf tubes. Whole kidneys were cut longitudinally and either snap frozen in liquid nitrogen for subsequent mRNA extraction or fixed in 10% neutral buffered formalin before embedding in paraffin for histological analysis.

### Assessment of renal function

Plasma samples were prepared from whole blood by centrifugation at 1000 x *g* for 5 minutes followed by aliquoting of the supernatant, snap freezing in liquid nitrogen and storage at −80°C until analysis without freeze-thaw. The plasma creatinine and albumin, urinary albumin and creatinine were measured as described previously (Mole et al., 2016). Urinary albumin excretion was expressed as albumin/creatinine ratio (ACR).

### Histology and digital image analysis

Histological sections (4 µm) of formalin-fixed paraffin embedded kidney tissue were de-waxed and taken through a decreasing series of graded alcohols to water. Hematoxylin and eosin (H&E) staining was performed according to standard protocols. The H&E stained sections were scored in a blinded fashion for assessment of tubular necrosis in the outer medulla (Wu et al., 2014). 10 representative random fields at a magnification of ×200 per section for each sample were examined. The percentage of tubules in the corticomedullary junction that displayed cellular necrosis and a loss of brush border were counted.

To assess the extent of apoptotic cell death induced by IRI, we performed terminal deoxynucleotidyltransferase dUTP-mediated nick end labeling (TUNEL) staining on paraffin-embedded kidney tissue sections using a commercially available kit (DeadEnd fluorometric TUNEL system; Promega, Madison, WI, USA). Briefly, formalin-fixed sections of 4 µm thickness were deparaffinized, hydrated, and incubated with 20 µg/mL proteinase K to strip proteins from the nuclei. Fragmented DNA was then identified by the incorporation of fluorescein-12-dUTP during an incubation step with terminal deoxynucleotidyltransferase at 37°C for 1 hour. Sections were stained by immersing the slides in 40 mL of propidium iodide solution freshly diluted to 1 µg/mL in PBS for 15 minutes. Microscopic images were acquired at ×10 magnification by using a Leica DC350F digital camera system equipped with Nikon Eclipse E800 Fluorescence microscope and Image-Pro Plus image analysis software (Media Cybernetics). Apoptotic cells (TUNEL-positive cells) were quantitatively assessed at ×100 magnification for 13 fields of tubular areas in a blinded manner using ImageJ as previously described (Mole et al., 2016).

Renal macrophages were identified by immunostaining for the tissue macrophage marker F4/80. Myeloperoxidase (MPO) positive cells were quantified in post-ischemic kidney sections as an index of neutrophil infiltration. The following primary antibodies were used: anti-mouse F4/80 monoclonal antibody (clone BM8) #14-4801 (eBioscience, Hatfield, UK) at 1:100 dilution and rabbit anti-MPO polyclonal antibody #1224 (Merck Millipore Corporation) at 1:1,000 dilution. Visualization was with diaminobenzoate (DAB) according to standard protocols. Type-specific control antibodies were used to distinguish background staining. Immunohistochemistry slides were scanned in their entirety using an Axio Scan.Z1 system (Zeiss microscopy GmbH, Oberkochen, Germany) and stored as .czi files before export as reduced-size .jpg files into ImageJ. Enumeration of F4/80^+^ and MPO^+^ cells was done using ImageJ as previously described (Mole et al., 2016), and expressed as positive cells per million pixels.

### RNA extraction and real time PCR

Total RNA was extracted from kidney tissue using an RNeasy Mini kit (Qiagen). 1µg of total RNA was used for first strand cDNA synthesis using a QuantiTect Reverse Transcription Kit (Qiagen). Expression of genes was determined by real-time PCR. Specific TaqMan primers and probes for kynurenine 3-monooxygenase (*Kmo*), kynureninase (*Kynu*), kynurenine aminotransferase (*Kat2*), 3-hydroxyanthranilic acid oxidase (*Haao*), interleukin-6 (*Il6*) and tumour necrosis factor α (*Tnfa*) were purchased from Life Biotechnologies. 18S ribosomal RNA was used as a reference gene. Amplification of cDNA samples was carried out using TaqMan® Fast Universal PCR Master Mix (AB Applied Biosystem) under the following conditions: 1 minute denaturation at 95°C, 45 cycles of 15 seconds at 95°C and 30 seconds at 60°C. Thermal cycling and fluorescence detection were conducted in a StepOne real time PCR system (Applied Biosystems). All reactions were carried out in triplicate and the cycle threshold (Ct) numbers of the target gene and reference gene in each sample were obtained. The mRNA levels of the target gene are presented as relative quantification (RQ) values.

### Liquid chromatography-tandem mass spectrometry (LC-MS/MS) analysis of kynurenine pathway metabolites

Samples of plasma and kidney tissue homogenate were diluted at a ratio of 2:5 in 5 mM ammonium formate containing 0.1% trifluoracetic acid. Protein was precipitated by the addition of ice-cold 100% trichloroacetic acid to samples, followed by incubation for 30 minutes at 4°C and centrifugation to obtain the supernatant. Serial dilutions of each metabolite were prepared over appropriate concentration ranges to prepare a calibration curve to permit quantitation. 10 µL volumes of each sample were injected onto a Waters Select HSS XP column (30 mm x 100 mm, 2.5 µm, Waters Corp, Elstree, Herts) using a Waters Acquity UPLC autosampler, coupled to an ABSciex QTRAP 5500 mass analyser. The flow rate was 0.35 mL/min at 25°C. Separation was carried out using a water:methanol gradient (both containing 0.1% formic acid). Conditions were 50:50 water:methanol to 40:60 over 60 seconds, 40:60 to 35:65 over 180 seconds, hold 35:65 for 110 seconds, 35:65 to 50:50 over 10 seconds and re-equilibration at 50:50 for 200 seconds. The total run time was 10 minutes. The mass spectrometer was operated in positive electrospray mode. The transitions for the protonated analytes were kynurenine, m/z 209-192; 3-hydroxykynurenine, m/z 225-202; tryptophan, m/z 205-188; kynurenic acid, m/z 190-144; and 3-hydroxyantranillic acid, m/z 154-136. Collision energies were 29, 15, 11, 31 and 33 eV respectively. Data was acquired and processed using analyst quantitation software (ABI Sciex).

### Experimental design and Statistical analysis

A simple 2 x 2 factorial design was used to compare experimental IRI versus a sham procedure in *Kmo*^null^ and *Kmo*^wt^ mouse strains. Group sizes were determined by a prospective power calculation using G-power(tm) (Faul et al., 2007) using input parameters from previous IRI experiments using the same IRI method and background mouse strain. A detectable effect size of 0.80 with power 1-β = 0.80 and significance, α=0.05, resulted in group sizes of n=6 mice per group. All data are expressed as mean ± SEM. All data were subjected to a one-sample Kolmogorov-Smirnov test to check whether data adhered to a Normal distribution. Normally-distributed data were analysed by one-way ANOVA followed by Tukey’s multiple comparison test. Data that did not follow a Normal distribution were analysed by Kruskal–Wallis test. All statistical analyses were performed using GraphPad(tm) Prism version 6.0d for Macintosh (GraphPad Software, San Diego, CA).

## AUTHOR CONTRIBUTIONS

Conceptualization, X.Z. and D.J.M.; Methodology, X.Z.; A.Z.; M.B.; K.M. and D.J.M; Investigation, X.Z.; A.Z.; M.B. and D.J.M.; Writing – Original Draft, X.Z. and D.J.M.; Writing – Review & Editing, X.Z.; K.M.; S.P.W.; J.H.; S.E.M.H. and D.J.M.; Funding Acquisition, D.J.M.; Supervision, D.J.M.; X.Z.; S.P.W. and S.E.M.H.; All authors read, discussed, and agreed with the final version of this manuscript.

## ACKNOWLEDGMENTS

We thank Spike Clay, Gary Borthwick, all members of the Biological Research Facility; Forbes Howie; Paul Fitch; Melanie McMillian, Debbie Mauchline and the SURF Facility; for their invaluable technical and/or management assistance. This work was supported by a Clinician Scientist Fellowship from the Health Foundation/Academy of Medical Sciences to DJM. DJM now acknowledges the support of the MRC through a Senior Clinical Fellowship.

